# Metal-induced energy transfer uncovers activation-induced axial reorganization of signaling complexes inside cells

**DOI:** 10.64898/2026.04.23.719849

**Authors:** Eike Wienbeuker, Arthur Felker, Oleksii Nevskyi, Steffen T. Harms, Daniel Marx, Kevin Tanzusch, Anna Chizhik, Rainer Kurre, Changjiang You, Jörg Enderlein, Jacob Piehler

**Affiliations:** Department of Biology/Chemistry and Center for Cellular Nanoanalytics, Osnabrück University, Barbarastr. 11, Germany; Third Institute of Physics (Biophysics), Georg August University, Friedrich-Hund-Platz 1, Germany

**Keywords:** Cytokine receptor, JAK-STAT signaling, intrinsically-disordered region, metal-induced energy transfer, surface nanopatterning, fluorescence lifetime imaging

## Abstract

Transmembrane signaling mediated by cytokine receptors orchestrates key cellular processes such as proliferation, differentiation, and immune responses. While numerous high-resolution structures of cytokine receptor ectodomains are available, the structural organization of the largely disordered intracellular domain (ICD) has remained unclear. Here, we interrogate the axial organization of cytokine receptor signaling complexes at the plasma membrane by metal-induced energy transfer (MIET). For this purpose, we leveraged biofunctionalized nanodot arrays (bNDAs) to capture cell surface receptors at a defined distance from the substrate. Readout by fluorescence lifetime imaging microscopy enabled quantifying axial distances of proteins in the plasma membrane of cells at both ensemble and single-molecule levels with a resolution of ∼1 nm. Using the prototypic, biomedically relevant class I cytokine receptor GP130 as a model system, we uncover by MIET that the ICD extends randomly into the cytosol in the resting state, but surprisingly undergoes an axial compaction upon signal activation. These results demonstrate the potential of bNDA-supported MIET for resolving the axial architecture of signaling complexes within the cellular context.

## Introduction

Multicellular organisms rely on elaborate intercellular communication networks in which a multitude of biochemical messengers is recognized by cognate receptors at the cell surface. Binding of messengers to cell surface receptors initiates intricate structural rearrangements that propagate signals across the plasma membrane, thereby orchestrating fundamental cellular processes including proliferation, differentiation, and immune responses (*1-3*). Different types of receptors utilize distinct mechanisms of signal propagation. For G protein-coupled receptors (GPCRs), ligand-induced conformational changes of the transmembrane helices have been characterized at atomic resolution by crystallography and cryo-electron microscopy (cryo-EM), establishing a paradigm for transmembrane signal propagation (*4, 5*). In contrast, the molecular principles governing downstream signal activation by single-pass transmembrane receptors, such as class I/II cytokine receptors (CRs), remain far less understood. Although high-resolution cryo-EM structures of several CR ectodomain (ECD) complexes have been reported (*6-8*), the intracellular domains (ICDs) of these receptors have remained structurally unresolved due to their inherent conformational heterogeneity.

CRs are activated by ligand-induced formation of dimeric or higher-order oligomeric assemblies (*9*), which triggers structural reorganization of the largely disordered ICDs (*10, 11*). Transient recruitment of effector proteins leads to the formation of dynamic signaling complexes at the plasma membrane. Yet how intrinsically disordered ICDs are spatially organized relative to the membrane, and how this organization changes upon signal activation, remains poorly understood. Current insight into these processes derives from molecular dynamics simulations and *in vitro* experiments, which often lack the complexity of the native cellular environment (*12*). Consequently, the axial architecture of CR signaling complexes at the plasma membrane has remained inaccessible. This impedes mechanistic understanding of the critical activation events at the plasma membrane for downstream signaling.

Here, we report a methodology to quantify the axial organization of CR signaling complexes at the plasma membrane with nanometer resolution by metal-induced energy transfer (MIET) (*13-15*). MIET leverages distance-dependent plasmonic fluorescence quenching near a metal surface to report axial positions with a dynamic range of ∼100 nm. As a model system, we employed glycoprotein 130 (GP130, encoded by *IL6ST*), a prototypic class I CR that is activated by IL-6 presented via its co-receptor IL-6Rα (**Fig. 1**A), as well as diverse other cytokines and their respective co-receptors (*16*).

**Fig. 1.**
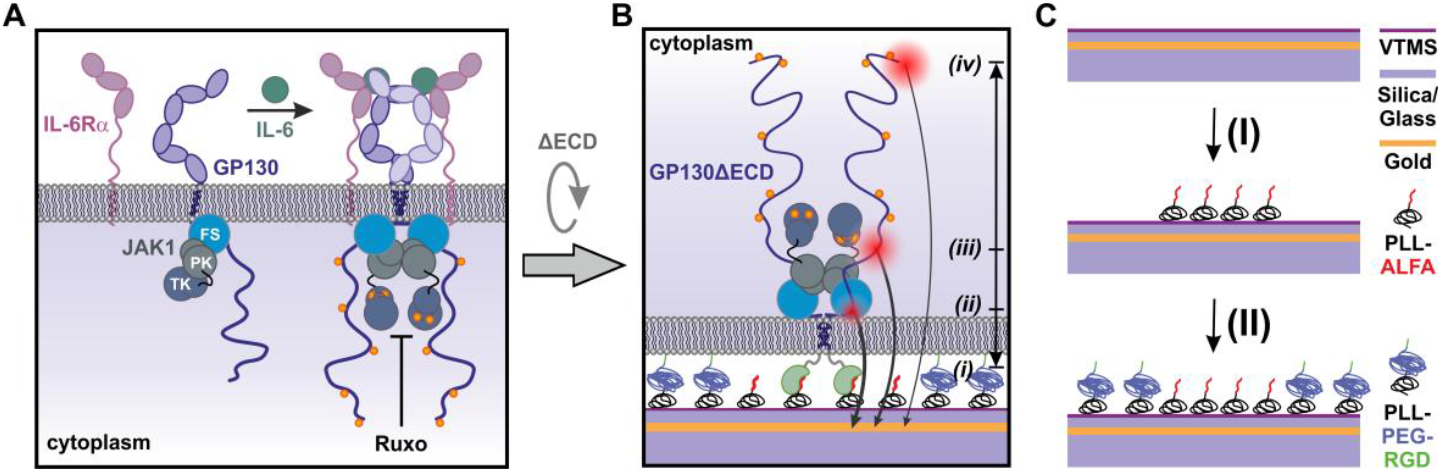
Probing the axial organization of GP130 signaling complexes at the plasma membrane. **(A)** Cell surface GP130 is dimerized by its agonist IL-6, which is presented via the co-receptor IL-6Rα. GP130 dimerization activates JAK1, associated with GP130 via its FERM-SH2 (FS) domain, by enabling interactions between the regulatory pseudokinase (PK) domains that promote cross-phosphorylation of the tyrosine kinase (TK) domains. Activated JAK1 subsequently phosphorylates several Tyr residues in the C-terminal IDR of GP130 (pTyr, orange). JAK1 activity can be blocked by the inhibitor Ruxolitinib (Ruxo). **(B)** Strategy for MIET-based axial distance mapping of the GP130 ICD. GP130, with its ECD replaced by an ALFA nanobody, is captured into ALFA-tag functionalized nanodots, positioning the ICD at a well-defined distance from the gold surface. Fluorophores at defined positions along the ICD (ii-iv) report axial distances via MIET-dependent fluorescence lifetime quenching. **(C)** Fabrication of ALFA-functionalized bNDAs on MIET substrates in two steps: Capillary nanostamping of PLL-ALFA onto a VTMS-silanized MIET substrate coated with a silica spacer (I), followed by backfilling with PLL-PEG-RGD (II).

GP130 is ubiquitously expressed across almost all human tissues and cell types, with important functions in coordinating immune responses, inflammation, cardiogenesis, liver regeneration and diverse other physiological processes. Agonist-induced GP130 dimerization activates downstream signaling via the cytosolic Janus family tyrosine kinase (JAK) member JAK1, which non-covalently associates with the membrane-proximal box 1/box 2 motifs of GP130 and is activated by cross-phosphorylation within dimers (*17-19*). Activated JAK1 phosphorylates five tyrosine residues within the ∼250-amino acid C-terminal intrinsically disordered region (IDR), generating docking sites for effector and regulator proteins including signal transducer and activator of transcription (STAT) proteins, most prominently STAT3, as well as diverse other effector proteins (**Fig. 1**A) (*20-22*). Dysregulation of the GP130-STAT3 signaling axis is implicated in diverse inflammatory diseases and cancer (*23*), and therefore emerged as an increasingly promising target for pharmaceutical intervention (*24-26*).

Aiming to resolve the axial conformational organization of the GP130 IDR by MIET, we devised biofunctionalized nanodot arrays (*27, 28*) (bNDAs) for capturing GP130 at a well-defined distance from the substrate surface (**Fig. 1**, B and C). By replacing the receptor’s ECD with a nanobody for immobilization into bNDAs, this strategy positions signaling complexes in close proximity to the gold layer while locally concentrating them within individual nanodots to enhance measurement contrast. Through systematic truncations in the ICD (**Fig. 1**B, positions ii-iv), we mapped the axial architecture of GP130 with nanometer resolution by fluorescence lifetime imaging microscopy (FLIM) at both ensemble and single-molecule levels. In the presence of the JAK inhibitor Ruxolitinib (Ruxo), our measurements surprisingly indicate that activation of downstream signaling compacts the C-terminal IDR, which can be explained by altered interactions of the tyrosine residues with active JAK1. These results establish MIET in bNDAs as a broadly applicable methodology to systematically resolve the axial organization and dynamics of signaling complexes at the plasma membrane in their native cellular context.

## Results

### Ultracompact NDA surface architecture for tight coupling with cellular systems

Precise MIET measurements of intracellular signaling complexes at the plasma membrane require their positioning at a well-defined and reproducible distance from the substrate surface. We addressed this by employing spatially resolved surface functionalization by capillary nanostamping (*27, 29*) of poly-L-lysine conjugated with the ALFAtag (*30*) (PLL-ALFA) onto microscopy slides silanized with vinyltrimethoxysilane (VTMS) to optimize surface wetting (*28*). For MIET experiments, microscopy substrates were coated with a 10 nm gold layer and a silica spacer (*31*) of defined thickness depending on the demands in different experiments (**Fig. 1**C), yielding arrays of ALFAtag-functionalized nanodots suitable for capturing proteins fused to an ALFA-tag nanobody (ALFAnb). *In vitro* characterization was performed by incubating bNDAs with ALFAnb fused to a monomeric enhanced green fluorescent protein (mEGFP-ALFAnb), followed by staining with ATTO643-conjugated anti-GFP nanobody enhancer (EN^ATTO643^, **Fig. 2**A) (*32*). Total internal reflection fluorescence (TIRF) microscopy confirmed effective binding of mEGFP-ALFAnb and EN^ATTO643^ into bNDAs (**Fig. 2**B and fig. S1), yet reliable quantification of nanodot diameters and contrast was not possible by diffraction-limited imaging (*28*). We therefore turned to DNA-PAINT super-resolution microscopy (*33*), which revealed densely packed, high-contrast nanodots with diameters below 500 nm (fig. S1).

**Fig. 2.**
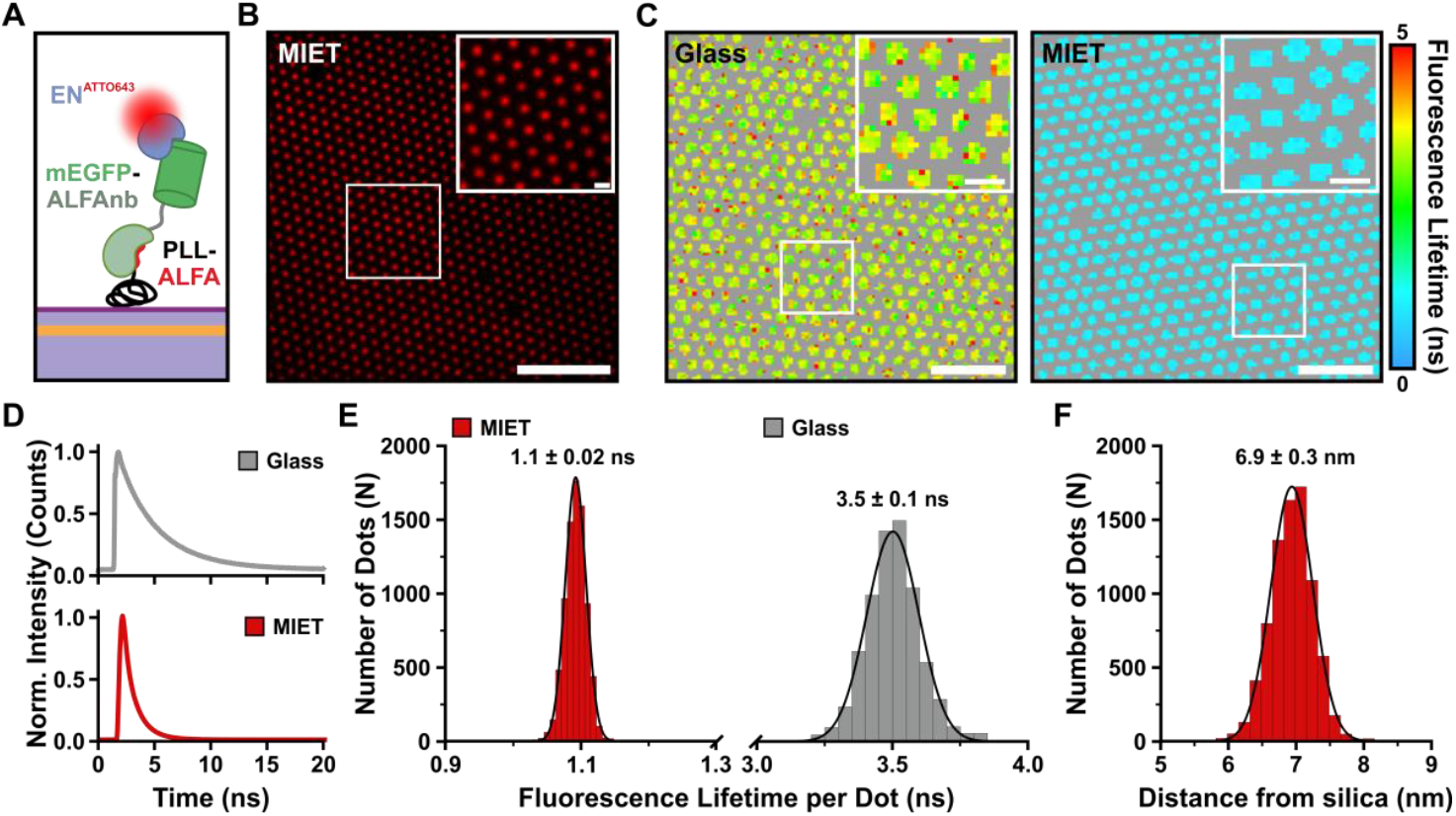
bNDAs establish a uniform molecular platform for MIET. **(A)** Schematic of the surface functionalization strategy: mEGFP-ALFAnb captured into PLL-ALFA bNDAs is stained with EN^ATTO643^ for MIET-based axial distance measurements. The MIET substrate comprised a 30 nm silica spacer. **(B)** Representative TIRF microscopy image of EN^ATTO643^-stained bNDAs on a MIET substrate. Inset shows a magnified view of the marked region. Scale bars: 10 µm; inset: 1 µm. **(C)** Representative fluorescence lifetime images of EN^ATTO643^ on glass (left) and MIET substrates (right). Insets show magnified views of the marked region. Scale bars: 5 µm; insets: 1 µm. **(D)** Representative normalized fluorescence decay curves of ATTO643 on glass (top) and MIET (bottom) substrates. **(E)** Per-nanodot fluorescence lifetime distributions on MIET (red; *n* = 8098 nanodots) and glass substrates (grey; *n* = 6975 nanodots). Solid lines represent Gaussian fits. **(F)** Axial distance distribution of ATTO643 from the silica surface (*n* = 8098 nanodots). Solid line represents Gaussian fit.

We next assessed MIET-induced lifetime quenching of ATTO643 within bNDAs by FLIM via time-correlated single photon counting (TCSPC), comparing fluorescence lifetimes on MIET substrates with a 30 nm silica spacer to an unquenched state on glass. These measurements confirmed the expected strong lifetime quenching (**Fig. 2**, C and D), with per-nanodot lifetimes of 1.10 ± 0.02 ns and 3.50 ± 0.10 ns on MIET and glass substrates, respectively (**Fig. 2**E). Fluorescence lifetimes were extracted using an analysis workflow comprising bNDA localization, per-nanodot lifetime fitting, and quality filtering (fig. S2, details in Methods). Using a calculated MIET calibration curve of ATTO643 considering optical parameters and the experimentally determined lifetime on glass (fig. S2), a mean distance *d* = 6.9 ± 0.3 nm of the dye from the silica surface was obtained (**Fig. 2**F).

### NDAs achieve well-defined transmembrane architectures in cells

Previous MIET experiments have revealed highly variable distances between the basal plasma membrane and the substrate surface (20-70 nm) (*13, 34*). With the additional top silica layer (20-30 nm) on the gold film, such conditions dictate the dynamic range for precise quantification of axial distances from the membrane. Capturing transmembrane proteins into bNDAs directly addresses this limitation by positioning them at a defined and reproducible proximity to the substrate. We therefore devised a model transmembrane protein with fluorescent reporters on both sides of the plasma membrane to validate this approach by dual-color MIET measurements. This construct comprised an extracellular ALFAnb-mEGFP and a cytosolic HaloTag separated by a transmembrane helix (ALFAnb-mEGFP-TMD-HaloTag), which is targeted into the plasma membrane via an N-terminal signal peptide, positioning the reporters at axial distances *d*_1_ and *d*_2_ from the substrate (**Fig. 3**A). HeLa cells expressing ALFAnb-mEGFP-TMD-HaloTag were cultured on PLL-ALFA NDAs and subsequently stained with HTL-JFX549, confirming nanopatterned reorganization of the construct in both the mEGFP and JFX549 fluorescence channels (**Fig. 3**B). To quantify the axial distances, cells were fixed, stained with EN^ATTO643^, and fluorescence lifetimes of both organic dyes were measured. Single-nanodot lifetime analysis revealed strong quenching within nanodots on MIET substrates relative to glass for both ATTO643 and JFX549 (**Fig. 3**, C and D and fig. S3, A and B). Per-nanodot lifetime analysis yielded 0.61 ± 0.04 ns and 1.74 ± 0.10 ns for ATTO643 and JFX549 on MIET substrates (**Fig. 3**E), which are substantially reduced compared to the unquenched reference lifetimes of 3.66 ± 0.13 ns and 3.79 ± 0.16 ns on glass, respectively (fig. S3, C and D). We calculated dye-specific MIET calibration curves (**Fig. 3**F) and obtained axial distances of *d*_1_ = 6.7 ± 0.7 nm and *d*_2_ = 20.3 ± 1.4 nm from the silica surface, yielding a *Δd* of 13.6 nm between the reporter molecules (**Fig. 3**G).

**Fig. 3.**
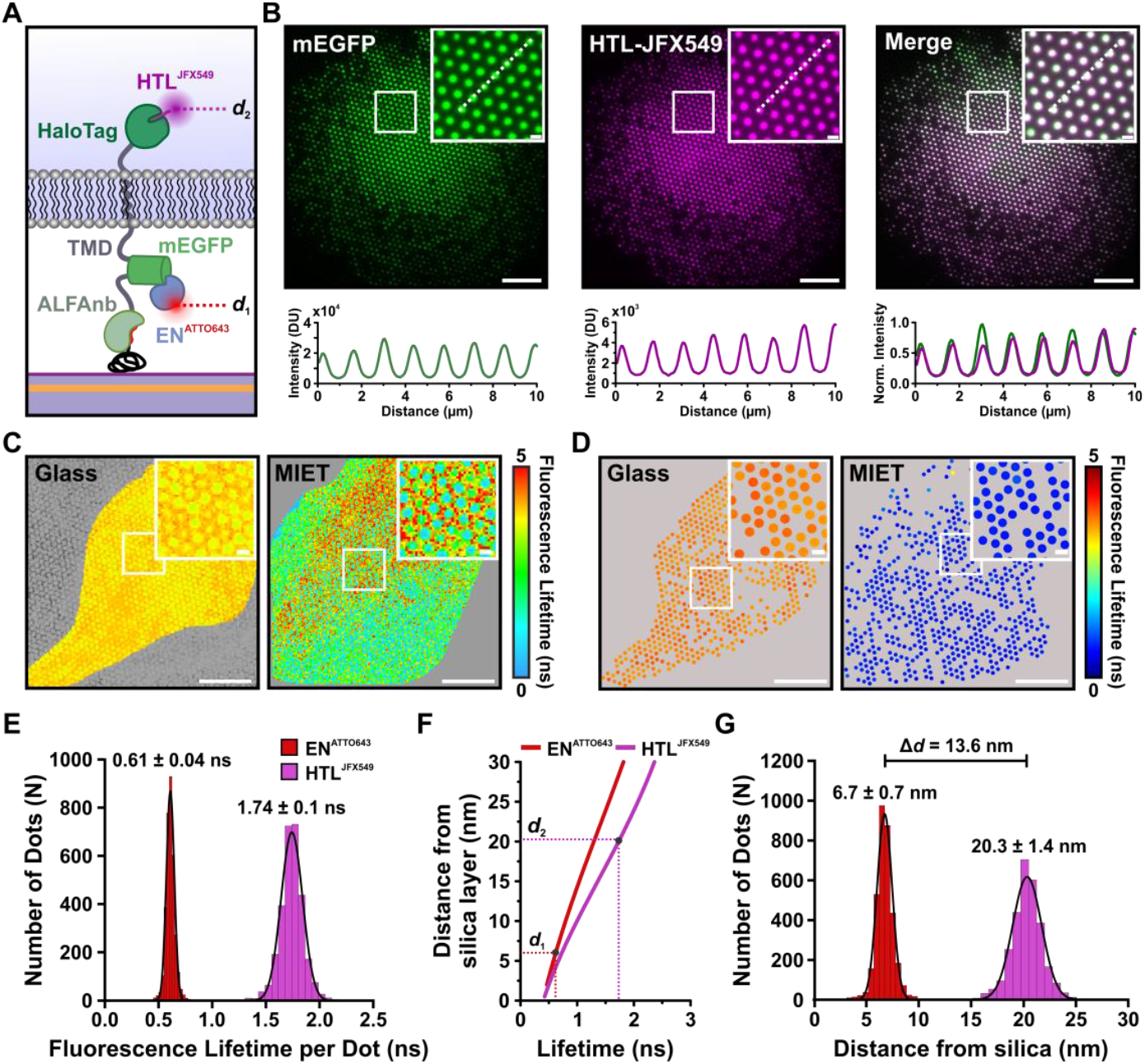
Architecture of a model transmembrane protein resolved by dual-color MIET in cells. **(A)** Schematic of the model transmembrane construct ALFAnb-mEGFP-TMD-HaloTag captured into PLL-ALFA NDAs via its extracellular ALFAnb. The extracellular mEGFP stained with EN^ATTO643^ and the cytosolic HaloTag labeled with HTL-JFX549 report axial distances *d*_1_ and *d*_2_ from the substrate surface, respectively. **(B)** Representative dual-color TIRF microscopy images showing mEGFP, HTL-JFX549, and merged fluorescence channels with corresponding intensity line profiles along the indicated white dashed lines. Insets show magnified views of the marked region. Scale bars: 10 µm; insets: 1 µm. **(C)** Representative raw fluorescence lifetime images of EN^ATTO643^ on glass and MIET substrates (20 nm silica spacer). Insets show magnified views of the marked region. Scale bars: 10 µm; insets: 1 µm. **(D)** Representative results from single-nanodot fluorescence lifetime analysis of the cell shown in C on glass and MIET substrates. Insets show magnified views of the marked region. Scale bars: 10 µm; insets: 1 µm. **(E)** Per-nanodot fluorescence lifetime distributions on MIET substrates for EN^ATTO643^ (red; *n* = 3428 nanodots, 9 cells) and HTL-JFX549 (magenta; *n* = 3022 nanodots, 7 cells). Solid lines represent Gaussian fits. **(F)** MIET calibration curves for EN^ATTO643^ (red) and HTL-JFX549 (magenta). Dashed lines indicate the axial distances *d*_1_ and *d*_2_ corresponding to the measured lifetimes. **(G)** Axial distance distributions of EN^ATTO643^ (*d*_1_, red) and HTL-JFX549 (*d*_2_, magenta) from the silica surface. Solid lines represent Gaussian fits. *Δd* marks the axial separation of both fluorescent reporters across the plasma membrane.

This value is in good agreement with structural modeling, predicting a *Δd* of 10.1-14.2 nm between the EN^ATTO643^ and HTL-JFX549 labeling sites by considering the flexibility of the linker connecting the TMD to the cytosolic HaloTag (fig. S4, A and B). These results confirm that capture into bNDAs effectively pulls the ectodomain of the model transmembrane protein towards the surface, establishing the desired transmembrane architecture for quantitative MIET measurements in cells.

### Axial mapping of the GP130 ICD reveals random coil extension in the resting state

We next aimed to interrogate the axial organization of the GP130 ICD in its resting state. To maintain a well-defined transmembrane architecture while positioning the ICD in close proximity to the MIET substrate, we replaced the N-terminal ectodomain of GP130, which comprises >800 amino acids, but is only necessary for ligand-induced homo- and heterodimerization (*6, 35*), with an ALFAnb for capturing into bNDAs. A C-terminal mEGFP was used for quantifying the distance from the substrate surface, yielding ALFAnb-GP130ΔECD-mEGFP (**Fig. 4**A). This construct was co-expressed with the FERM-SH2 domain of JAK1 (JAK1(FS)-mScarlet), which binds to the membrane-proximal box 1/box 2 motifs of GP130, thereby ensuring structural organization of the cytosolic receptor domain. Under these conditions, signal activation is effectively suppressed as both the pseudokinase and kinase domains are absent (*36*). As reference points for the axial interrogation, we systematically truncated the receptor’s ICD and generated three constructs: one lacking the entire cytoplasmic domain (ΔICD), one truncated after the JAK1-binding motifs (Box1/2), and one comprising the full-length ICD (f-ICD), enabling quantification of *d*_1_ and *d*_2_ as schematically depicted in **Fig. 4**A. HeLa cells co-expressing ALFAnb-GP130ΔECD-mEGFP and JAK1(FS)-mScarlet were cultured on PLL-ALFA NDAs, followed by fixation, permeabilization, and staining mEGFP with EN^ATTO643^ for fluorescence lifetime quantification. TIRF microscopy confirmed effective co-capturing of receptor and JAK1(FS) into nanodots and successful post-fixation staining (**Fig. 4**B). MIET was then quantified at ensemble (Ens.) level by analyzing single-nanodot fluorescence lifetimes on glass and MIET substrates for all three GP130 constructs (**Fig. 4**C; fig. S5, A to C; and table S1). Furthermore, we applied FLIM-based DNA-PAINT (*37*) (FL-PAINT) to quantify axial distances at the single-molecule level (*15*), using an anti-GFP nanobody conjugated to a docking strand for staining with a complementary ATTO643-conjugated imager strand (**Fig. 4**D; fig. S5, A to C; and table S2).

**Fig. 4.**
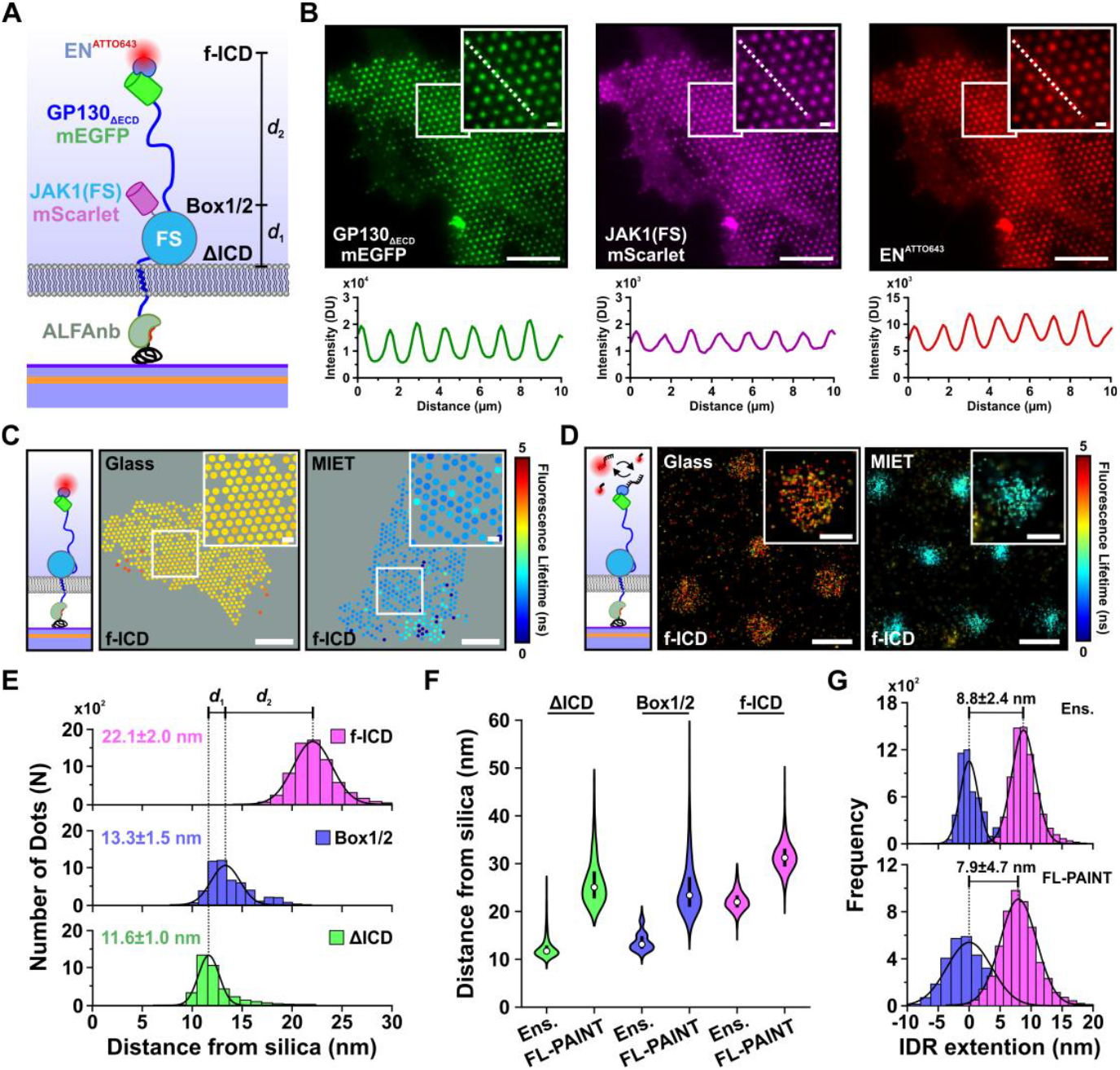
MIET resolves random coil extension of the GP130 IDR in the resting state. **(A)** Schematic of ALFAnb-GP130ΔECD-mEGFP captured into PLL-ALFA NDAs, co-expressed with JAK1(FS)-mScarlet. Three ICD truncation constructs (ΔICD, Box1/2, and f-ICD) enable quantification of *d*_1_ and *d*_2_. **(B)** Representative TIRF microscopy images showing GP130ΔECD-mEGFP and JAK1(FS)-mScarlet, with mEGFP stained post-fixation with EN^ATTO643^, and corresponding intensity line profiles along the indicated white dashed lines. Insets show magnified views of the marked region. Scale bars: 10 µm; insets: 1 µm. **(C)** Representative single-nanodot fluorescence lifetime analysis of EN^ATTO643^ on glass and MIET substrates (20 nm silica spacer) for the f-ICD construct. Insets show magnified views of the marked region. Scale bars: 10 µm; insets: 1 µm. **(D)** Representative FLIM-based DNA-PAINT (FL-PAINT) images of the f-ICD construct on glass and MIET substrates. Insets show magnified views of an individual nanodot. Scale bars: 1 µm; insets: 200 nm. **(E)** Axial distance distributions from ensemble single-nanodot analysis for ΔICD (green; *n* = 4002 nanodots, 8 cells), Box1/2 (blue; *n* = 5123 nanodots, 7 cells), and f-ICD (magenta; *n* = 6956 nanodots, 8 cells). Dashed lines indicate *d*_1_ and *d*_2_. Solid lines represent Gaussian fits. **(F)** Comparison of axial distances obtained by ensemble and FL-PAINT analysis for ΔICD (green), Box1/2 (blue), and f-ICD (magenta). *n* values and summary statistics are listed in tables S1 and S2. **(G)** Extension of the GP130 IDR quantified by ensemble (top) and FL-PAINT (bottom) analysis. Solid lines represent Gaussian fits.

The axial distance distributions obtained from ensemble single-nanodot analysis revealed distinct architectures for all three GP130 variants (**Fig. 4**E): a small but significant separation of *d*_1_ = 1.7 nm from the cytosolic membrane leaflet (ΔICD, 11.6 ± 1.0 nm) to the JAK1-binding motif (Box1/2, 13.3 ± 1.5 nm), and an extension of *d*_2_ = 8.8 nm of the downstream IDR to the C-terminus of GP130 (f-ICD, 22.1 ± 2.0 nm). Single-molecule analysis overall yielded larger axial distances (**Fig. 4**F and tables S1 and S2), which may be caused by non-random orientation (*13, 34*), due to electrostatic repulsion of the docking strand from the negatively charged cytosolic surface of the plasma membrane, and further amplified upon hybridization with the imager strand. Nevertheless, referencing f-ICD distances to the Box1/2 position resulted in a consistent IDR extension (Ens.: 8.8 ± 2.4 nm; PAINT: 7.9 ± 4.7 nm), confirming the robustness of the measurement across both approaches (**Fig. 4**G). The absolute *d* _1_ values, however, did not reveal the expected axial distance between the ΔICD and Box1/2 constructs in PAINT measurements, which we ascribed to bias related to oligonucleotide labeling. Therefore, ensemble analysis was prioritized in subsequent experiments. The observed IDR extension is quantitatively in line with a random coil conformation: assuming an effective backbone segment length of 0.36-0.38 nm per residue, the ∼220 amino acid IDR corresponds to a Flory exponent of 0.58-0.59, characteristic of typical disordered chains in buffer solution (*38, 39*). Collectively, these results demonstrate that the GP130 C-terminal IDR extends freely into the cytosol in the resting state with a total axial distance of 10.5 nm (*d*_1_ + *d*_2_), rather than adopting a membrane-associated conformation as previously proposed for related receptors based on *in vitro* and *in silico* data (*10, 40*).

### JAK1 activation induces axial rearrangement of the GP130 IDR

GP130 activation drives JAK1-mediated phosphorylation of five IDR tyrosines, which serve as docking sites for STATs and other effector proteins (*41, 42*). Whether this process alters the axial organization of the disordered chain, however, remained unresolved. We therefore captured GP130/JAK1 signaling complexes in bNDAs to quantify activation-dependent axial changes by MIET (**Fig. 5**A). Interestingly, ALFAnb-GP130ΔECD-mEGFP co-expressed with JAK1-HT exhibited substantial ligand-independent STAT3 phosphorylation (fig. S6A). This constitutive activation was leveraged to control JAK1 kinase activity in GP130 bNDAs by the highly effective, non-toxic and reversible type I JAK inhibitor Ruxolitinib (Ruxo) (*43*).

**Fig. 5.**
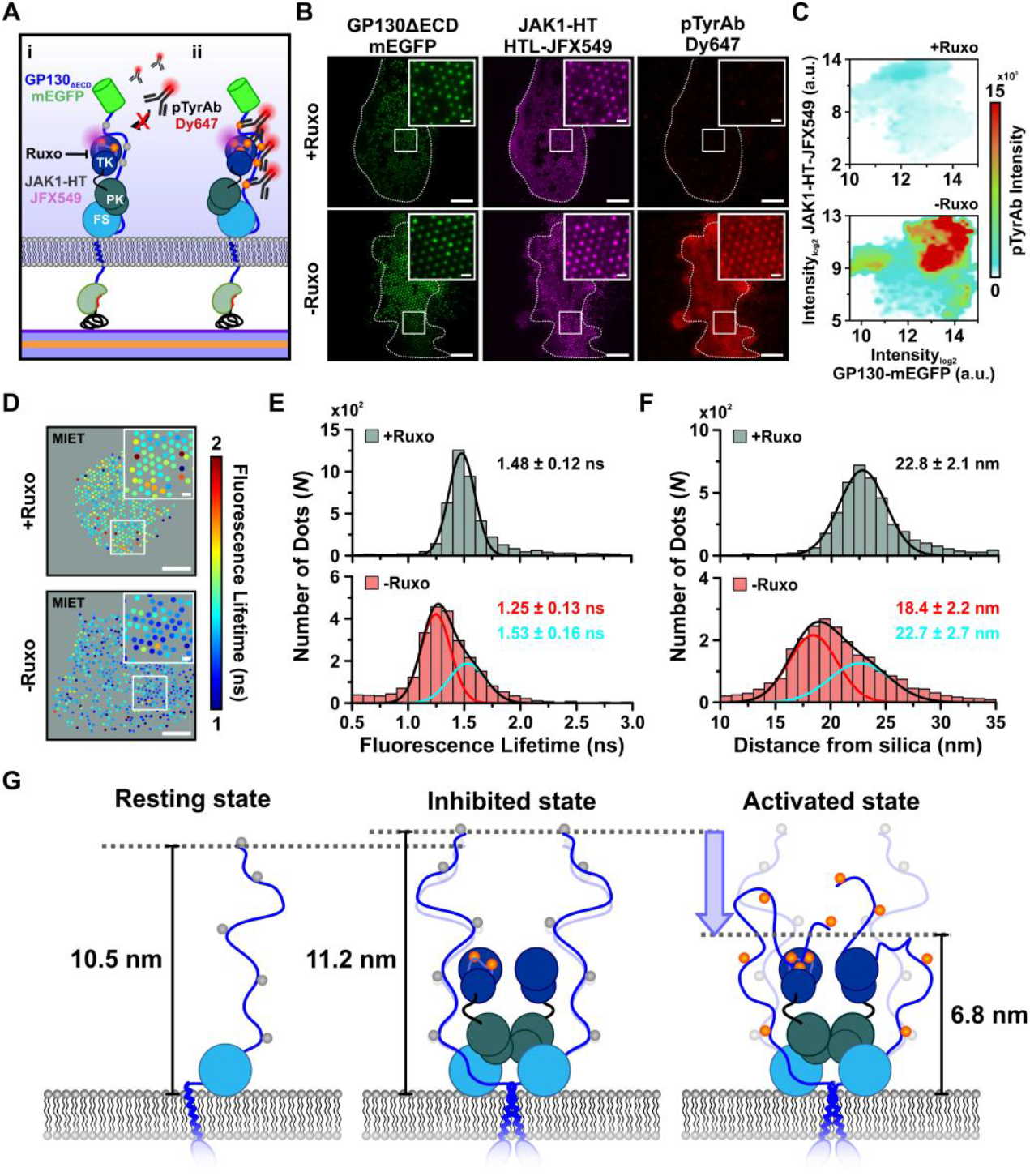
Activity-dependent axial reorganization of the GP130 IDR. **(A)** Schematic of the experimental design for probing activation-dependent axial changes of the GP130 IDR in bNDAs. ALFAnb-GP130ΔECD-mEGFP and JAK1-HT labeled with HTL-JFX549 are co-recruited into PLL-ALFA bNDAs, with kinase activity controlled by Ruxo (i) and tyrosine phosphorylation (pTyr, orange) detected with an anti-pTyr antibody (pTyrAb-Dy647) (ii). **(B)** Representative TIRF microscopy images showing ALFAnb-GP130ΔECD-mEGFP, JAK1-HT labeled with HTL-JFX549, and pTyrAb-Dy647 staining in the presence (+Ruxo, top) and absence (-Ruxo, bottom) of Ruxolitinib. Insets show magnified views of the marked regions. Scale bars: 10 µm; insets: 1 µm. **(C)** Single-nanodot correlation analysis of GP130ΔECD-mEGFP and JAK1-HT (HTL-JFX549) fluorescence intensities, color-coded by pTyrAb-Dy647 intensity, for +Ruxo (top; *n* = 7729 nanodots, 15 cells) and -Ruxo (bottom; *n* = 8174 nanodots, 21 cells) conditions. **(D)** Representative results from single-nanodot fluorescence lifetime analysis of EN^ATTO643^ on MIET substrates (20 nm silica spacer) in the presence (+Ruxo, top) and absence (-Ruxo, bottom) of the inhibitor. Insets show magnified views of the marked regions. Scale bars: 10 µm; insets: 1 µm. **(E)** Per-nanodot fluorescence lifetime distributions under +Ruxo (grey; *n* = 4460 nanodots, 9 cells) and -Ruxo (red; *n* = 2689 nanodots, 5 cells) conditions. Solid lines represent Gaussian fits. **(F)** Axial distance distributions for +Ruxo (grey) and -Ruxo (red) conditions, calculated from the fluorescence lifetimes shown in (E). Solid lines represent Gaussian fits. **(G)** Model of the three axial states of the GP130 IDR: the resting state with bound JAK1(FS) (10.5 nm), the Ruxo-inhibited state with full-length JAK1 (11.2 nm), and the activated state (6.8 nm). Distances refer to the IDR C-terminus above the inner plasma membrane leaflet.

Immunostaining with an anti-phosphotyrosine antibody (pTyrAb) of HeLa cells cultured on bNDAs confirmed robust, nanodot-selective Tyr phosphorylation activity upon washing out Ruxo. By contrast, negligible phosphorylation was observed upon JAK1 inhibition by Ruxo, confirming the capability to effectively control JAK1 activation by this inhibitor (**Fig. 5**, B and C, and fig. S7, A to C). These results confirm bNDA capturing of GP130:JAK1 complexes in a signaling-active, i.e. dimeric, state.

To quantify possible signaling-induced axial changes, fluorescence lifetimes of EN^ATTO643^ bound to the C-terminal mEGFP of ALFAnb-GP130ΔECD-mEGFP were measured on glass and MIET substrates in the presence and absence of Ruxo. In the presence of Ruxo (+Ruxo), a unimodal lifetime distribution centered at 1.48 ± 0.12 ns yielded a distance of 22.8 ± 2.1 nm (**Fig. 5**, D to F). Strikingly, heterogeneous lifetimes across individual nanodots were observed after washout of Ruxo (-Ruxo, **Fig. 5**D), which was reflected in a substantially broadened lifetime distribution, suggesting a signaling-related axial rearrangement of the GP130 IDR (**Fig. 5**E). Two distinct lifetime components centered at 1.25 ± 0.13 ns (64 %) and 1.53 ± 0.16 ns (36 %) were resolved, corresponding to axial distances of 18.4 ± 2.2 nm and 22.7 ± 2.7 nm, respectively (**Fig. 5**, E and F). The minor component was consistent with the Ruxo-treated, signaling-incompetent state, while the dominant component revealed a substantial decrease in axial distance. The bimodal character likely reflects heterogeneous JAK1 activation across individual nanodots, yielding a mixed population of activated and inactive complexes. Referencing these distances to the cytosolic membrane position established by the ΔICD construct places the IDR C-terminus at ∼11.2 nm and ∼6.8 nm above the membrane leaflet in the inhibited and activated states, respectively (**Fig. 5**G). The former is consistent with the freely extended conformation obtained for the resting state in the presence of JAK1-FS (10.5 nm), while the latter indicates a 4.4 nm compaction during signaling. This demonstrates that kinase activity rather than JAK1 binding per se drives the axial rearrangement of the GP130 IDR. Together, these results establish that bNDA-supported MIET resolves activation-induced changes in the axial organization of intrinsically disordered receptor domains within the cellular context.

## Discussion

Cytokine receptor signaling depends on structural rearrangements that propagate from the extracellular side of the receptor to its cytosolic face, yet the molecular organization of the predominantly intrinsically disordered intracellular domains that orchestrate these responses has remained largely unexplored. In contrast to the well-defined structures of extracellular and transmembrane entities (*6, 44*), IDRs are far more dynamic, transiently engaging effector proteins and membrane surfaces (*12, 45, 46*). While small-angle x-ray scattering (SAXS), nuclear magnetic resonance (NMR), and molecular dynamics simulations have provided fundamental insights into the dynamic organization of such domains in well-defined molecular context (*10*), dedicated tools are required to validate *in vitro* predictions in the native cellular environment.

Here, we established bNDA-supported MIET as a generic approach for quantifying the axial organization of receptor signaling complexes at the plasma membrane with nanometer resolution directly in cells. Capturing transmembrane receptors into bNDAs offered two key advantages: (i) positioning the signaling complexes in close and well-defined proximity to the gold surface, within an optimal sensitivity range for MIET, and (ii) locally concentrating complexes within individual nanodots to substantially enhance signal-to-background ratio. Through optimization of the surface functionalization and capturing system, the distance between the plasma membrane and the substrate surface was reduced to ∼6-7 nm, and by adjusting the silica spacer thickness, the MIET sensitivity range can be further tuned to match the expected axial position of any given target. Together with single-nanodot lifetime analysis, this enabled nanometer axial discrimination measurement directly within cells.

The class I CR GP130 with its broad biological and biomedical relevance served as a model system for studying the axial organization of intrinsically disordered signaling complexes. The mechanisms underlying the broad functional plasticity and pleiotropy of GP130 family cytokines, which compromise successful pharmaceutical targeting, have largely remained enigmatic. While structural studies and live-cell single-molecule imaging suggest the formation of receptor dimers with structurally well-defined ectodomains (*6, 41, 47-50*), functional plasticity is most likely encoded in the conformationally flexible IDR with its multiple docking sites for diverse effector and negative regulator proteins. Mapping distinct axial positions along the cytosolic receptor domain by MIET revealed that the ∼220 amino acid C-terminal IDR freely extends ∼ 8.8 nm beyond the JAK1-binding motifs into the cytosol. Previous *in vitro* and *in silico* studies on a related class I cytokine receptor have suggested association of the ICD with the surface of the inner plasma membrane leaflet, which has been interpreted to potentially impose lateral constraints on IDR mobility and pre-organize phosphorylation sites near the membrane surface (*10*). By contrast, our studies suggest that the C-terminal IDR of GP130 freely extends into the cytosol as expected for a random coil, thereby ensuring that docking motifs are accessible to cytosolic effector proteins. The presence of full-length JAK1 in its inhibited state did not alter the axial position of the IDR compared to the resting state, which was mimicked by JAK1 lacking the PK-TK domains (**Fig. 5**G, center and left). Binding of Ruxo into the ATP-binding pocket of the JAK TK domain has been shown to induce a substantial conformational change, leading to a tyrosine-phosphorylated activation-loop that is fixed in an inaccessible conformation (*51-53*). We therefore assume that Ruxo-inhibited full-length JAK1 does not promote additional interactions with the GP130 ICD as compared to the truncated JAK1, in line with a negligible difference in the axial organization. However, interactions between the JAK1 PK domains should induce substantial GP130 dimerization, which may be responsible for a slight increase in distance upon bringing the IDRs of the two GP130 into closer proximity (**Fig. 5**G). Most surprisingly, however, a pronounced reorganization of the disordered chain was observed, resulting in a compaction of 4.4 nm upon GP130 activation. We expected that Tyr phosphorylation of the ICDs, which was confirmed by antibody staining, rather leads to extension of the IDR due to effector binding to these docking sites. This effect can be ascribed to interactions of Tyr residues in the GP130 IDR with the membrane-proximal JAK1 TK substrate binding site as depicted in **Fig. 5**G, right. However, docking of STATs (and potentially other effector proteins), which form dimers already in the resting state (*54*), may crosslink docking sites in the IDRs and thus cause the observed compaction.

These proof-of-concept experiments establish the potential of this new methodology to robustly assess axial conformations of intrinsically disordered receptor domains directly in the crowded and chemically complex cytoplasmic environment, opening new perspectives for dissecting the dynamic interplay between multiple components within intact signaling complexes. While the present study focused on a single timepoint of activation, future experiments resolving the temporal dynamics of IDR reorganization during signaling and termination will be essential for a complete mechanistic picture. Correlating distances of co-recruited effector and regulator proteins such as STATs and SOCS would directly link axial geometry to local signaling output. Realizing these possibilities will, however, require methodological advances. Quantifying MIET in living cells will be essential, as fixation-induced conformational artifacts cannot be entirely excluded. FL-PAINT experiments already indicated that the staining procedure itself may affect MIET measurements, further motivating the implementation of reversible labeling strategies compatible with live-cell imaging (*55-57*). Especially single-molecule approaches could not only enable mapping of conformational heterogeneity, but also capture the dynamics of rapid fluctuations between states. Beyond lifetime-based measurements, intensity-based readouts could provide complementary insights into conformational dynamics at higher temporal resolution.

With its generic concept, the methodology can be readily adapted to other receptor families with large cytoplasmic disordered regions, including receptor tyrosine kinases, the T cell receptor, or Wnt co-receptors, which have been shown to locally induce condensation of their effector proteins at the membrane (*27, 58-61*). Compared to Förster resonance energy transfer (FRET), MIET offers a ∼10-fold larger dynamic range, covering the full conformational space of disordered receptor domains and their signaling complexes. Importantly, no fluorescence acceptor is required, and therefore associated labeling stoichiometry artifacts are excluded. Taken together, we have established a methodology for interrogating the axial architecture of signaling complexes in their native cellular environment, addressing a fundamental gap in our ability to decode the structural logic of transmembrane signaling at the plasma membrane.

## Materials and Methods

### Preparation of MIET substrates

MIET substrates were fabricated on glass coverslips by sequentially depositing layers of 2 nm titanium, 10 nm gold, 1 nm titanium, and 20 or 30 nm silicon dioxide. The layers were deposited by electron-beam evaporation using an electron beam source (Univex 350, Leybold) under high-vacuum conditions (∼10^−6^ mbars). The thin silicon dioxide spacer layer was included to achieve an optimal distance between the sample and the gold layer, corresponding to the most sensitive region of the MIET curve.

### Synthesis of PLL-ALFAtag

PLL-ALFAtag was synthesized through a two-step coupling reaction. Briefly, cysteine-containing peptide (Ac-CPSRLEEELRRRLTE; 1.4 µmol) was dissolved in 100 µL HEPES-buffered saline (HBS; 20 mM HEPES, 150 mM NaCl, pH 7.5) followed by addition of MAL-PEG_2_-NHS (1.7 µmol) dissolved in 100 µL HBS. After 15 min of gentle shaking at room temperature, a solution of 3.5 mg poly-L-lysine hydrobromide (molecular weight: 15 - 30 kDa) in 400 µL HBS was added to this tube. The conjugation reaction was carried out for 12 h at room temperature (25°C) with continuous stirring. The reaction mixture was purified by dialysis (14 kDa molecular-weight cut-off) against 2 L Milli-Q water at 4°C for 24 h. The final product was obtained by lyophilization of the dialysate, yielding a white powder.

### Capillary nanostamping

Mesoporous silica stamps were prepared according to previously established protocols (*29, 62*). Post-synthesis, the silica stamps were neutralized and washed with Milli-Q water at 60°C for 24 h to remove residual surfactants and chemical contaminants. Solvent exchange was performed by incubating the stamps in ethanol for 24 h, with the solvent refreshed at least three times. The equilibrated stamps were stored for one week prior to application. Microscopy substrates were plasma-cleaned (Plasma Cleaner Femto 1A, Diener Electronic GmbH) for 10 min (glass coverslips, ϕ24 mm / thickness 170 µm from Marienfeld-Superior) or 5 min (MIET substrates). Afterwards, the substrates were immersed in 4 % (v/v) vinyltrimethoxysilane in toluene and incubated for 18 h at room temperature. Prior to stamping, the substrates were rinsed with toluene and ethanol to remove excess silane and solvent. Finally, the substrates were gently blown dry with nitrogen gas. Immediately before the stamping procedure, silica stamps were equilibrated in Milli-Q water for 2 h. After incubating with 20 µL of PLL-ALFAtag (1 mg/mL, dissolved in HBS: 20 mM HEPES and 150 mM NaCl, pH 7.5) for 15 min, excess ink was removed via a gentle stream of nitrogen gas. Ink-loaded stamps were mounted onto a stamp holder using double-sided adhesive tape and brought into contact with silanized microscopy substrates. Following the stamping procedure, substrates were passivated to minimize non-specific protein adsorption on non-patterned regions by incubation with 1 mg/mL solution of PLL-PEG-Methoxy for 45 min. In experiments involving live cells, the passivation was performed using a 50:50 (v/v) mixture of PLL-PEG-Methoxy and PLL-PEG-RGD (*63*). Finally, the patterned substrates were washed with Milli-Q water and stored at 4°C until further experimental use.

### Cell culture and transfection

The plasmids used for cell transfection are listed in table S3. HeLa cells (obtained from German Collection of Microorganisms and Cell Cultures, ID: ACC57) were maintained in MEM supplemented with fetal bovine serum, non-essential amino acids, and HEPES (MEM++) at 37°C and 5 % CO_2_. Cells were passaged at ∼70 % confluency by trypsin-ethylenediamine tetra-acetic acid (EDTA) treatment and split at a 1:6 ratio. Cells seeded on 60 mm dishes were transiently transfected using a polyethylenimine (PEI) transfection protocol: 3 µg DNA plasmid was mixed with 12 µL PEI in 300 µL Milli-Q water containing 150 mM NaCl for 15 min at room temperature, and applied to the cells for transfection. After incubation in MEM++ for 8 h at 37°C, cells were washed three times with warm PBS and cultured in fresh MEM++ for 12 h for protein expression and cell regeneration.

### Ruxolitinib treatment, fixation and antibody staining

Transfected HeLa cells were treated with Trypsin/EDTA, seeded on NDA-functionalized substrates, and incubated for 2 h at 37°C and 5 % CO_2_. The cells were then treated with 20 µM Ruxolitinib (Ruxo) for 2.5 h to inhibit JAK1 and thus suppress signaling. JAK activation was initiated by washing three times with 1 mL PBS to remove Ruxo, followed by incubation for 30 min at 37°C during which the HaloTag was stained with 20 nM HTL-JFX549. Control samples were also stained with 20 nM HTL-JFX549 for 30 min at 37°C, but remained in Ruxo-supplemented medium throughout. Cells were placed in SDS/ethanol-cleaned custom-built microscopy chambers and fixed with 4 % paraformaldehyde (PFA) (15 min, room temperature) and washed three times with 1 mL PBS. Simultaneous permeabilization and blocking were performed with 3 % bovine serum albumin (BSA) and 0.25 % Triton X-100 supplemented with anti-pTyr primary antibody (mouse) overnight at 4°C. After washing three times with PBS containing 0.5 % BSA (PBSA), samples were incubated with Dy647-conjugated anti-mouse secondary antibody for 4 h at room temperature, washed three times with 1 mL PBSA, and stored in HBS until imaging.

### Total internal reflection fluorescence microscopy (TIRFM)

TIRF imaging was performed at 25°C on an inverted microscope (Olympus IX-83) equipped with a 4-line TIRF condenser (cellTIRF (MITICO), Olympus) and a 100× oil immersion objective (UPLAPO100XOHR, NA 1.5, Olympus). mEGFP, JFX549, and ATTO643/Dy647 were excited at 488 nm (LuxX 488-200, Omicron), 561 nm (2RU-VFL-P-500-561-B1R, MPB Communications), and 642 nm (2RU-VFL-P-500-642-B1R, MPB Communications), respectively. Fluorescence was collected through a TIRF pentaband polychroic beamsplitter (Semrock zt405/488/561/640/730rpc) and channel-specific bandpass filters: 525/35 (mEGFP), 600/50 ET (JFX549), and 685/50 ET (ATTO643/Dy647). Channels were acquired sequentially with an sCMOS camera (Hamamatsu ORCA-FusionBT). Focus was stabilized by a hardware autofocus system (IX3-ZDC2, 830 nm, Olympus). Image acquisition was performed using 2×2 pixel binning (effective pixel size: 130 nm) and controlled via CellSens Dimension software (v3.1, Olympus).

### Fluorescence lifetime imaging (FLIM)

Confocal imaging and FLIM were performed on an inverted microscope (Olympus IX-81) equipped with a motorized stage (SCAN IM 112×74 mm, Märzhäuser) and a 60× oil immersion objective (UPLXAPO, NA 1.42, Olympus). Laser excitation at 488, 559, and 635 nm was provided by an Olympus laser box with AOTF combiner. Fluorescence was detected by three photomultiplier tubes (R7862, Hamamatsu) via a two-mirror galvanometer scanning unit at 512×512 pixel resolution (pixel size: 103 nm). The confocal pinhole was set to approximately one Airy unit. FLIM measurements were performed using a PicoQuant FLIM upgrade kit. Fluorescence lifetimes of JFX549 and ATTO643 were acquired by time-correlated single photon counting (TCSPC) using pulsed laser diodes at 560 nm (LDH-D-TA-560, 20 MHz) and 640 nm (LDH-D-C-640, 20 MHz), respectively. Emitted fluorescence was spectrally isolated with bandpass filters (Semrock BrightLine 600/30 for JFX549; 690/70 for ATTO643) and detected by two independent PMA Hybrid 40 detectors synchronized via a MultiHarp 150 TCSPC module.

### FL-PAINT setup

Fluorescence lifetime measurements were performed on a custom-built confocal setup. Fluorescence excitation was done with a 640 nm, 40 MHz pulsed diode laser (PDL 800-B driver with LDH-D-C-640 diode, PicoQuant). After passing through a cleanup filter (MaxDiode 640/8, Semrock), a quarter-wave plate converted the linearly polarized laser light into circularly polarized light. Subsequently, the laser beam was coupled into a single-mode fiber (PMC-460Si-3.0-NA012-3APC-150-P, Schäfter + Kirchhoff) with a fiber coupler (60SMS-1-4-RGBV-11-47, Schäfter + Kirchhoff). After the fiber, the output beam was collimated by an air objective (UPlanSApo 10×/0.40 NA, Olympus). An ultraflat quad-band dichroic mirror (ZT405/488/561/640rpc, Chroma) reflected the excitation light toward the microscope. After passing a laser scanning system (FLIMbee, PicoQuant), the light was sent into the custom side port of the microscope (IX73, Olympus). The three galvo mirrors of the scanning system deflected the beam while preserving the beam position in the back focal plane of the objective (UApo N 100×/1.49 NA oil, Olympus). Sample position was adjusted by using the manual *XY* stage of the microscope (IX73, Olympus) and a *z* piezo stage (Nano-ZL100, Mad City Labs). Fluorescence light was collected by the same objective and descanned by the scanning system. An achromatic lens (TTL180-A, Thorlabs) focused the descanned beam onto a pinhole (100 μm P100S, Thorlabs). Backscattered/back-reflected excitation laser light was blocked by a long-pass filter (635 LP Edge Basic, Semrock). After the pinhole, the emission light was collimated by a 100 mm lens. An additional band-pass filter (BrightLine HC 679/41, Semrock) was used for further rejection of scattered excitation light. Finally, the emission light was focused onto a single-photon avalanche diode (SPAD) detector (SPCM-AQRH, Excelitas) with an achromatic lens (AC254-030-A-ML, Thorlabs). Output signals of the photon detectors were recorded with a TCSPC module (HydraHarp 400, PicoQuant) that was synchronized by a trigger signal from the excitation laser. Images were acquired with the software SymPhoTime 64 (PicoQuant), which controlled both the TCSPC electronics and the scanner. Typically, samples were scanned with a virtual pixel size of 100 nm, a dwell time of 2.5 μs per pixel, and a TCSPC time resolution of 16 ps.

### Fluorescence lifetime analysis

Fluorescence lifetime data were processed using the TrackNTrace (TNT) software framework in MATLAB (*64*). Temporal stacks were integrated into a single frame for ensemble-averaged analysis. Emitters were localized by 2D Gaussian fitting followed by Maximum Likelihood Estimation (MLE) to extract position, amplitude, background, and PSF standard deviation. Fluorescence lifetimes were determined by biexponential fitting of decay curves (bounds: 0.2-5 ns), initialized with *τ*_*1*_ at 0.2 ns and *τ*_*2*_ at the reference lifetime of the respective dye (10 iterations). Localizations were filtered by PSF sigma and fit quality, followed by manual ROI definition to exclude extracellular background and non-specific signal.

For single-molecule imaging, images were reconstructed from the raw scan data by combining every 10 scans into a single frame. Localization was performed in TrackNTrace using the Cross-Correlation detection plugin with default settings, followed by refinement with the TNT Fitter plugin employing pixel-integrated Gaussian maximum-likelihood fitting. Localizations detected in adjacent frames and separated by less than 100 nm were linked into a track, and the final position was determined by refitting the summed image data from all frames assigned to that track.

For ensemble glass controls and *in vitro* MIET samples, the amplitude-weighted mean lifetime of the two biexponential components was used. For cellular MIET samples, only the shorter lifetime component was considered for distance calculations. In single-molecule imaging experiments, fluorescence lifetimes were determined by mono-exponential MLE fitting.

For each condition, MIET calibration curves were generated using a custom written MATLAB script, accounting for substrate architecture (refractive indices and thicknesses of Ti, Au, and SiO_2_ spacer thickness), fluorophore emission wavelength, and quantum yield (*QY*). The QY was corrected by the ratio of the free-space lifetime measured on glass (*LT*_*FS*_) to the reference lifetime (*LT*_*R*_):

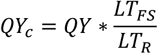

Distances were extracted by inverting the calibration curve and applying a linear fit to its dynamic range, converting per-nanodot lifetimes directly to axial distances. Calibration curves, lifetime histograms, and distance histograms were fitted in OriginPro 9.

### DNA-PAINT imaging and analysis

NDAs were functionalized with 200 nM mEGFP-ALFAnb in HBS for 15 min, washed five times with HBS, and incubated overnight at 4°C with 20 nM anti-GFP nanobody conjugated to docking strand F3 (Massive Photonics). Imaging was performed on the TIRF microscope described above at 25°C using 561 nm excitation in imaging buffer containing 50 pM Cy3B-conjugated imager strand F3 (Massive Photonics). Datasets consisted of 20,000 frames at 50 ms exposure with 2×2 pixel binning (effective pixel size: 130 nm); focus was maintained by hardware autofocus throughout. Raw data were analyzed using the Picasso software suite (*33*). Localizations were identified in Picasso Localize (box size: 7 px; net gradient threshold: 5,000-20,000) and filtered for localization precision (≤ 10 nm). Drift correction was performed by cross-correlation followed by fiducial marker refinement. Nanodot diameters were measured manually from localization boundaries; contrast was calculated as the ratio of mean nanodot to local background intensity.

### Flow cytometry

Antibodies used in the work are summarized in table S4. Transfected HeLa cells were detached by trypsin-EDTA treatment, pelleted (300 × g, 5 min), resuspended in MEM++, and seeded into a 96-well plate. HeLa WT and HeLa GP130/JAK1 double knockout (dKO) cells were treated with 20 nM Hyper-IL6 (HyIL6). Cells were fixed with 4 % PFA (15 min), washed with PBSA, pelleted (300 × g, 5 min), and permeabilized in ice-cold methanol (500 µL, 30 min). After washing with PBSA, cells were stained with 1 µL anti-pY705-STAT3-AF647 antibody for 1 h, washed twice with PBSA, and analyzed on a CytoFLEX flow cytometer (Beckman Coulter) at 60 µL/min. Instrument performance was verified using QC standardization beads (Beckman Coulter) according to the manufacturer’s instructions. From each sample, HeLa cells were selected based on characteristic scatter properties. Receptor-positive cells were gated for GP130 expression (FITC: 10^5^ to 50^5^; receptor low) and JAK1 co-expression (PB450: 10^4^-50^4^; kinase medium). Mean APC-A fluorescence intensity was extracted from this subpopulation, and the standard error of the mean (SEM) was calculated.

### Structural modeling of the mEGFP-TMD-HaloTag construct

A structural model of the ALFAnb-mEGFP-TMD-HaloTag construct was assembled in PyMOL (Schrödinger) to estimate axial inter-dye distances. The construct was arranged N-to C-terminally as: mEGFP-Linker 1 (GSGGS)-GP130 TMD-Linker 2 (GGSLEGHGTGSTGSGSS)-HaloTag7. The mEGFP-enhancer complex was modeled from PDB 3K1K; the HaloTag7-chloroalkane ligand (OEH) complex from PDB 6Y7A. The GP130 TMD was predicted by AlphaFold3 (pLDDT > 90) and aligned along the membrane normal at Z = 0. Extracellularly, the mEGFP C-terminus was positioned 1.9 nm from the TMD N-terminus, assuming full extension of Linker 1 (5 aa × 0.38 nm/aa). Intracellularly, two conformations of the 17-aa Linker 2 were modeled to capture its conformational range: a compact conformation defined by the root-mean-square end-to-end distance from a Worm-Like Chain (WLC) model:

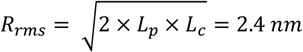

with a persistence length (L_p_) of 0.45 nm for GS-rich linkers (*65*) and a contour length (L_c_) of 6.46 nm (17 aa × 0.38 nm/aa), and a fully extended conformation at maximum contour length (L_c_ = 6.46 nm). Inter-dye distances were measured in PyMOL as the Z-coordinate difference between the C-terminal residue of the enhancer and the centroid of the HaloTag ligand. Structures were rendered in UCSF ChimeraX (*66*).

## Acknowledgements

We thank A. Budke-Gieseking, G. Hikade and W. Kohl (Osnabrück University) for technical support, and Janelia Materials for providing HTL-JFX549. This project was supported by the German Research Foundation (DFG) grants RTG2900 (no. 501879556) to J.P. and C.Y., SFB 1557 (no. 467522186, projects P13 and Z2) to J.P. and R.K., and by funding to J.E. from the European Research Council (ERC) under the European Union’s Horizon 2020 research and innovation program (AdR ERC Grant “smMIET” No. 884488).

## Author contributions

Conceptualization: E.W., A.F., C.Y. and J.P. Investigation: E.W. Methodology: E.W., A.F., S.H., K.T., A.C., R.K. and O.N. Data curation: E.W. Formal analysis: E.W. and A.F. Project administration: E.W., A.F. and J.P. Funding acquisition: J.P. and J.E. Resources: J.E. and J.P. Software: A.F., O.N., D.M. and J.E. Supervision: J.P. and J.E. Validation: E.W. and A.F. Visualization: E.W. and A.F. Writing – original draft: C.Y and J.P. Writing – review and editing: E.W., A.F., O.N., A.C., D.M., S.H., K.T., R.K., C.Y., J.E. and J.P.

## Data availability statement

All data needed to evaluate the conclusions in the paper are present in the paper or the Supplementary Materials. Further information and requests for resources and reagents should be directed to and will be fulfilled by the corresponding author on request.

