## Supplementary Information for "Metal-induced energy transfer uncovers activation-induced axial reorganization of signaling complexes inside cells"

**This PDF file includes:**

Figs. S1 to S7

Tables S1 to S5

### Supplementary Figures

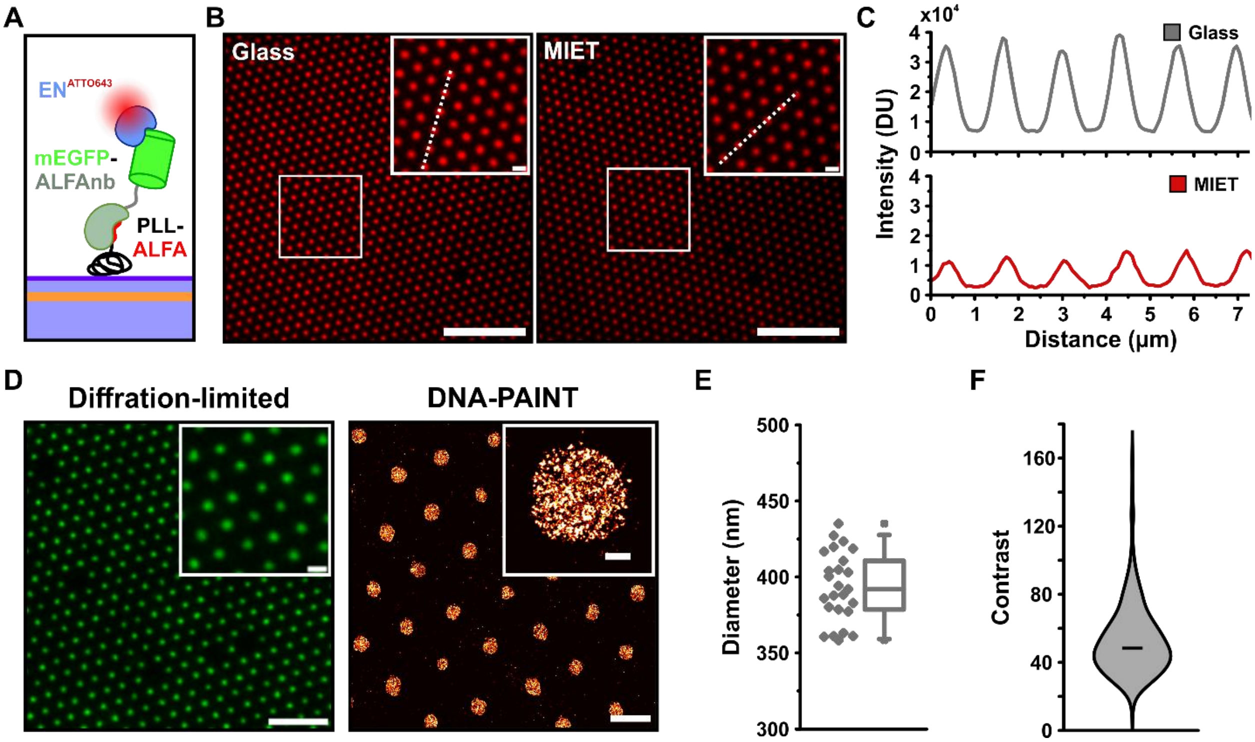


**Fig. S1. Characterization of PLL-ALFA bNDAs by TIRF microscopy and DNA-PAINT. (A)** Schematic illustration of the PLL-ALFA/ALFAnb-mEGFP/EN$^{ATTO643}$ biofunctionalization strategy. **(B)** Representative TIRF microscopy images of PLL-ALFA NDAs on glass (left) and MIET (right) substrates after staining_with EN$^{ATTO643}$. Dashed lines indicate the positions of intensity profiles shown in (C). Insets show magnified views. Scale bars: 10 µm; inset: 1 µm. **(C)** Fluorescence intensity profiles along the dashed lines shown in (B), demonstrating strong per-nanodot quenching on MIET substrates. **(D)** Diffraction-limited (left) and DNA-PAINT super-resolution (right) images of bNDAs after binding of mEGFP-ALFAnb. Insets show magnified views of individual nanodots. Scale bars for diffraction-limited image: 5 µm; inset: 1 µm. Scale bars for DNA-PAINT: 1 µm; inset: 100 nm. **(E)** Nanodot diameters quantified from DNA-PAINT images (*n* = 25 nanodots). **(F)** Per-nanodot contrast values determined from DNA-PAINT images (*n* = 291 nanodots).

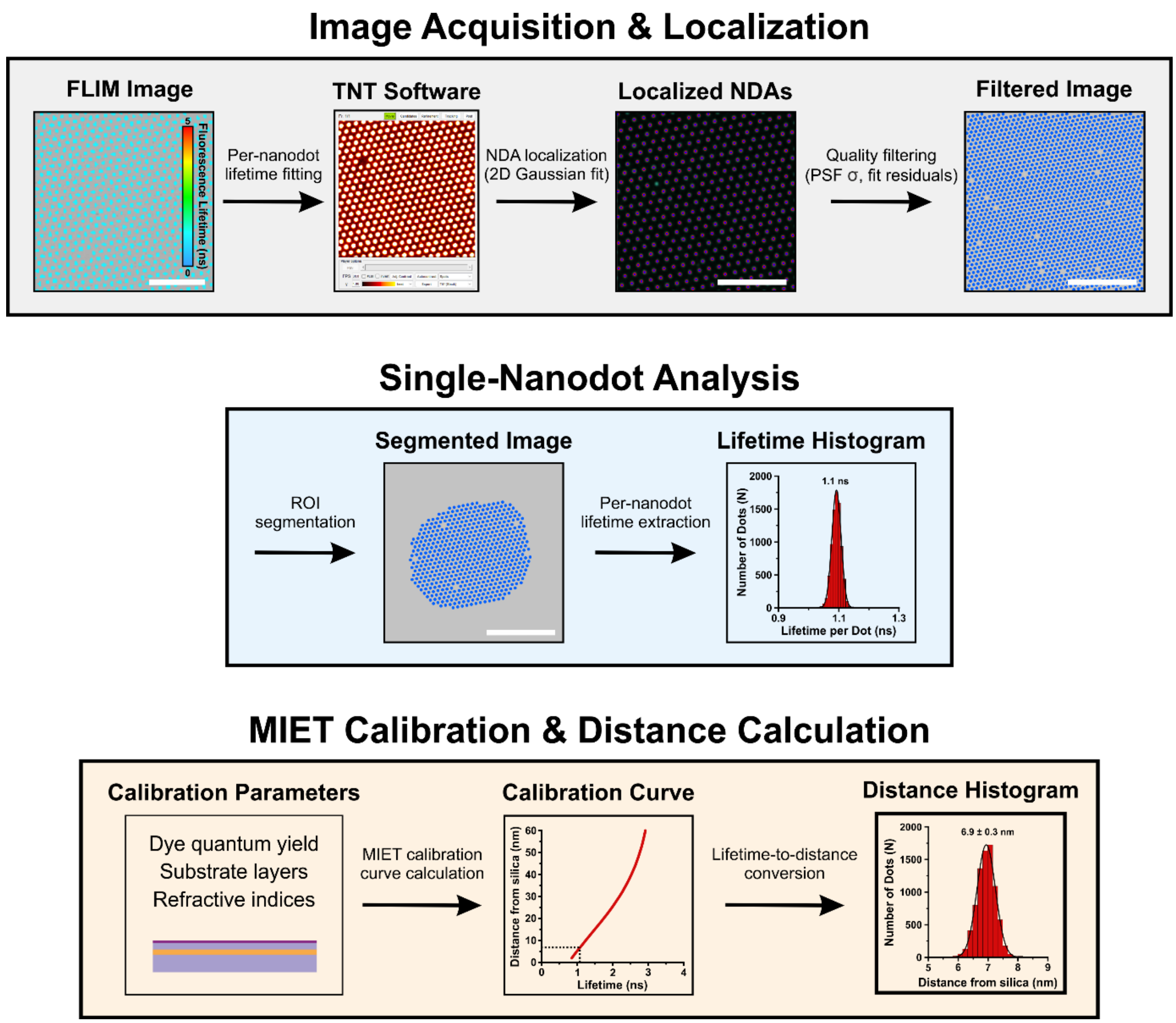


**Fig. S2. Single-nanodot fluorescence lifetime analysis workflow.** (Top) Image acquisition and localization workflow: Raw FLIM images are processed in TNT software through binning, bNDA localization, quality filtering, and per-nanodot lifetime fitting. (Middle) Single-nanodot analysis: ROI segmentation of the filtered image enables per-nanodot lifetime extraction, yielding a lifetime histogram. (Bottom) MIET calibration and distance calculation: a calibration curve is calculated from substrate layer thicknesses, refractive indices, dye quantum yield and the free-space lifetime on glass, and is used to convert per-nanodot lifetimes to axial distances from the silica surface.

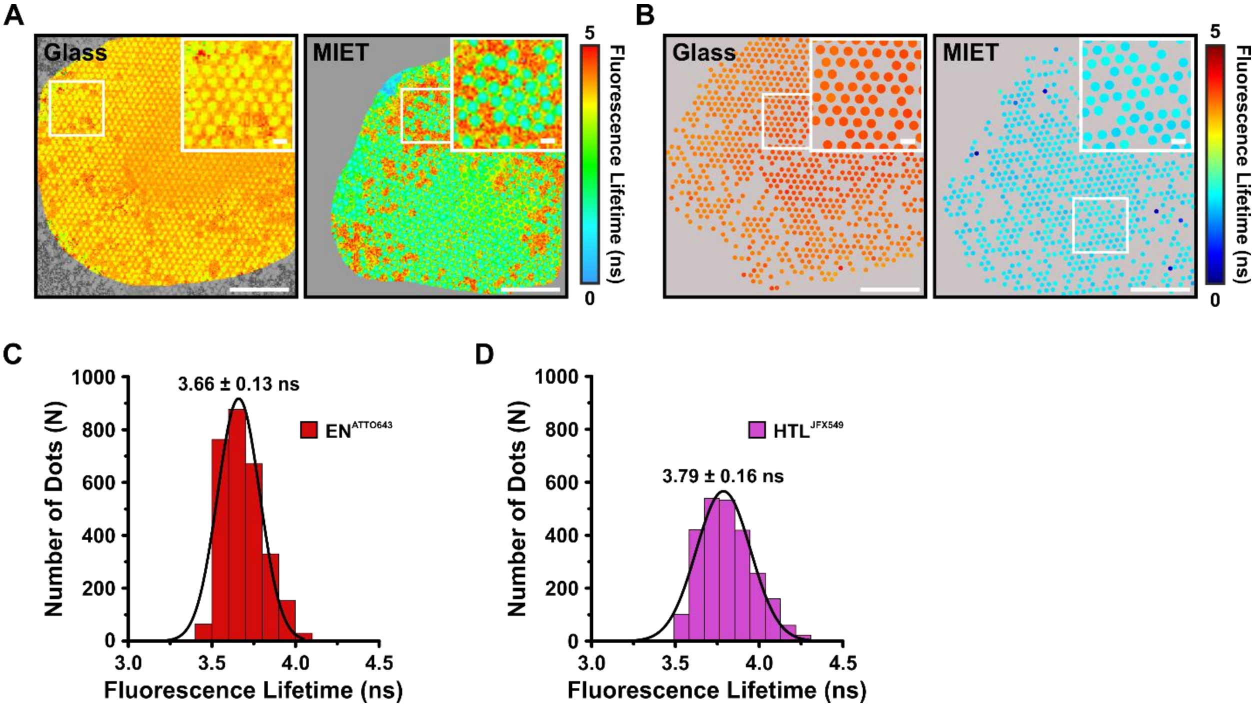


**Fig. S3. Complementary fluorescence lifetime data and quantification for HTL-JFX549 and EN$^{ATTO643}$ in bNDAs.** Complementary lifetime data for the experiment shown in Fig. 3. **(A)** Representative raw fluorescence lifetime images of HTL-JFX549 on glass and MIET substrates. Insets show magnified views of the marked region. Scale bars: 5 µm; insets: 1 µm. **(B)** Representative results from single-nanodot fluorescence lifetime analysis of the cell shown in A on glass and MIET substrates. Insets show magnified views of the marked region. Scale bars: 5 µm; insets: 1 µm. **(C, D)** Per-nanodot fluorescence lifetime distributions on glass substrates for EN$^{ATTO643}$ (red; 3.66 ± 0.13 ns; $n$ = 2889 nanodots, 5 cells) (C) and HTL-JFX549 (magenta; 3.79 ± 0.16 ns; $n$ = 2521 nanodots, 8 cells). Solid lines represent Gaussian fits.

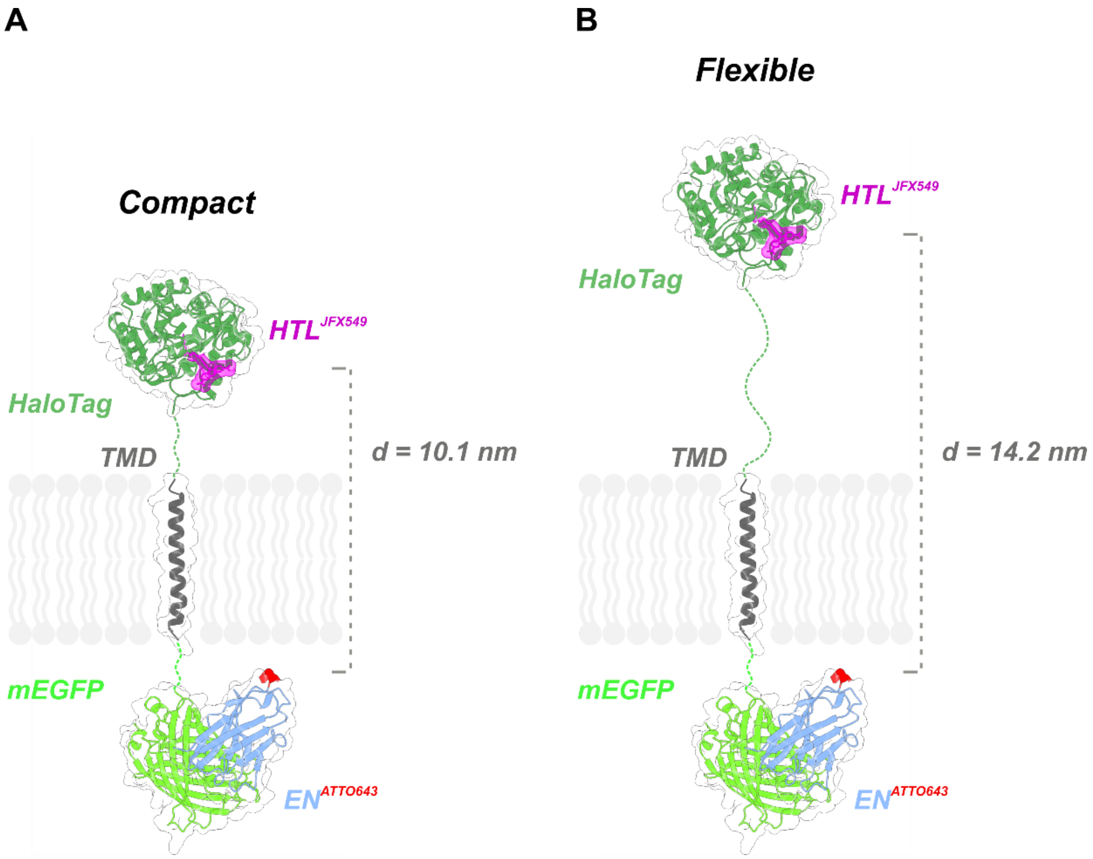


**Fig. S4. Structural modeling of labeling site distances in ALFAnb-mEGFP-TMD-HaloTag model transmembrane protein. (A, B)** Structural models in compact (**A**) and flexible (**B**) linker conformations. The extracellular mEGFP (light green) is stained with $EN^{ATTO643}$ (blue; ATTO643 in red), connected via the TMD (dark grey) to the cytosolic HaloTag (dark green) labeled with HTL-JFX549 (magenta). Flexible linkers are indicated by green dashed lines. The lipid bilayer is shown in grey. The axial distance *d* between the two labeling sites (A: 10.1 nm; B: 14.2 nm) defines the dynamic range expected from the flexibility of the intracellular linker. Protein models are based on PDB entries 3K1K (mEGFP-EN) and 6Y7A (HaloTag), combined with an AlphaFold3-predicted TMD. Further details are provided in the Methods section.

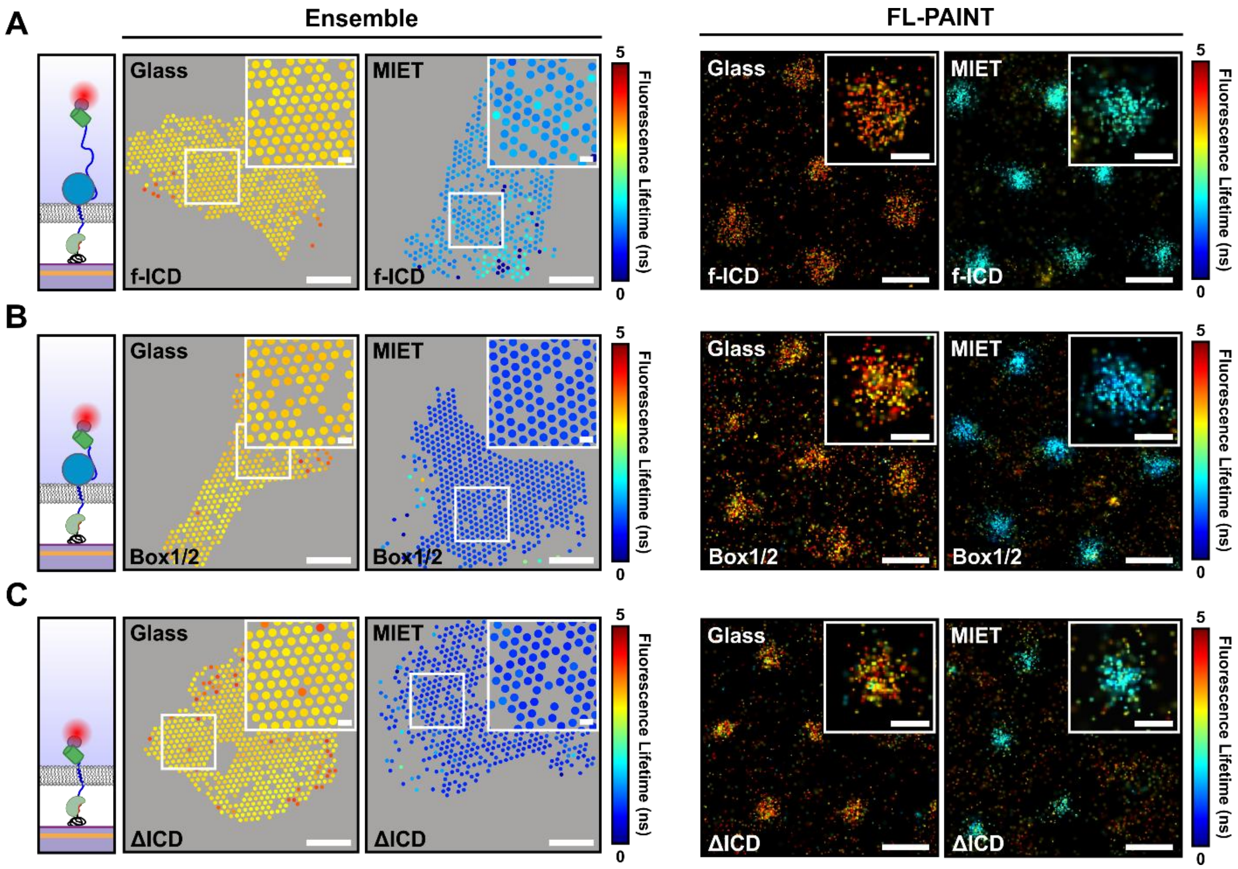


**Fig. S5. Single-nanodot ensemble FLIM and FL-PAINT images of all three GP130 constructs. (A-C)** Representative single-nanodot fluorescence lifetime images of $EN^{ATTO643}$ on glass and MIET substrates obtained by ensemble analysis (left) and FL-PAINT (right) for f-ICD (A), Box1/2 (B), and ΔICD (C). Cartoons illustrate the respective ICD truncation construct. Insets show magnified views of the marked region. Scale bars: 10 µm (ensemble); 1 µm (FL-PAINT); insets: 1 µm (ensemble); 200 nm (FL-PAINT).

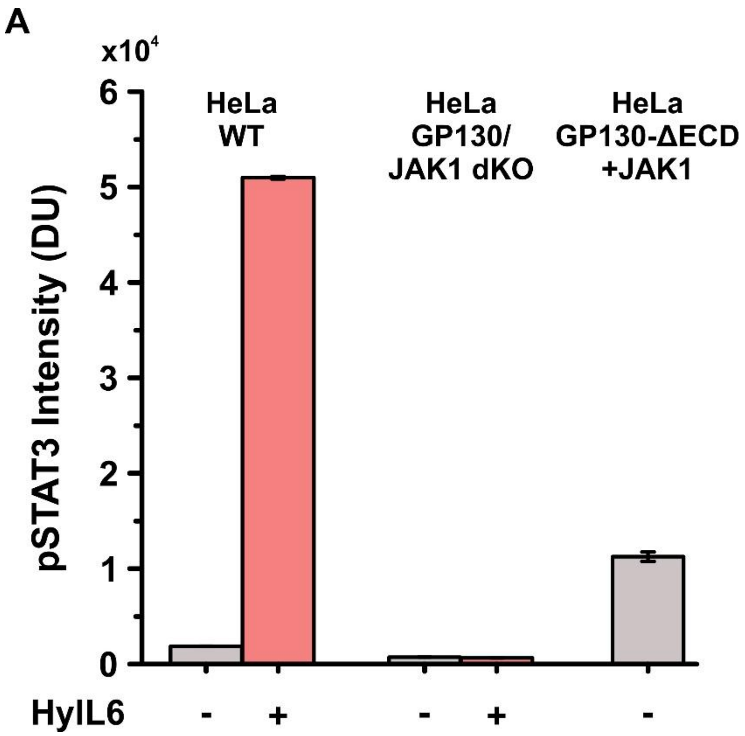


**Fig. S6. Ligand-independent STAT3 phosphorylation of GP130ΔECD co-expressed with JAK1. (A)** Flow cytometry quantification of pSTAT3 intensity in HeLa wild-type cells (-HyIL6: *n* = 35116 cells; +HyIL6: *n* = 52963 cells) and HeLa GP130/JAK1 double knockout (dKO) cells (-HyIL6: *n* = 163327 cells; +HyIL6: *n* = 209406 cells) without and with ligand stimulated by HyIL6, respectively, in comparison with HeLa cells expressing GP130ΔECD co-transfected with JAK1 in the absence of ligand (*n* = 1500 cells). Data are presented as mean ± SEM.

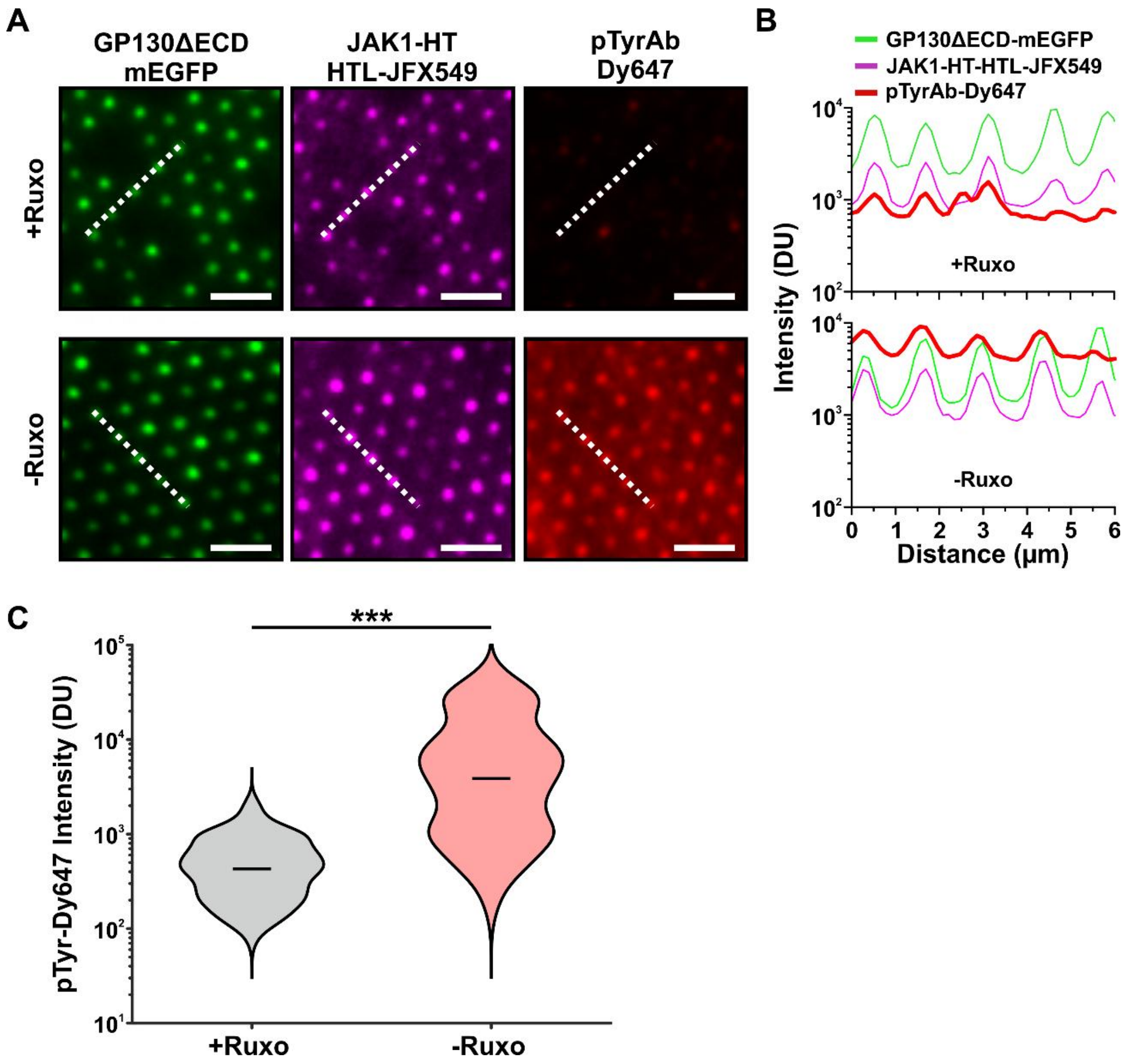


**Fig. S7. Phosphotyrosine staining of GP130/JAK1 signaling complexes in bNDAs. (A)** Representative TIRF microscopy images of HeLa cells co-expressing ALFAnb-GP130ΔECD-mEGFP and JAK1-HT labeled with HTL-JFX549, stained with a pTyrAb-Dy647, in the presence (+Ruxo) and absence (-Ruxo) of the JAK inhibitor Ruxolitinib. Scale bars: 2 µm. **(B)** Intensity line profiles of GP130ΔECD-mEGFP, JAK1-HT-HTL-JFX549, and pTyrAb-Dy647 along the dashed white lines indicated in (A) for +Ruxo (top) and -Ruxo (bottom) conditions. **(C)** Violin plot of nanodot-selective pTyrAb-Dy647 intensities under +Ruxo (*n* = 7729 nanodots, 15 cells) and -Ruxo (*n* = 8174 nanodots, 21 cells) conditions. Horizontal lines indicate medians. ***p < 0.001, two-sample Kolmogorov-Smirnov test.

### Supplementary Tables

#### Supplementary Table S1: Summary of Lifetimes and Distances from single-nanodot fluorescence lifetime analyses.

| Condition | Substrate | #Nanodots | Lifetime (ns)[1] | Distance (nm)[2] |
|---|---|---|---|---|
| mEGFP-ALFAnb (in vitro) | Glass | 6975 | 3.50±0.10 | 6.9±0.3 |
| | MIET | 8098 | 1.10±0.02 | |
| mEGFP-TMD-HaloTag (EN$^{ATTO643}$) | Glass | 2889 | 3.66±0.13 | 6.7±0.7 |
| | MIET | 3428 | 0.61±0.04 | |
| mEGFP-TMD-HaloTag (JFX549) | Glass | 2521 | 3.79±0.16 | 20.3±1.4 |
| | MIET | 3022 | 1.74±0.10 | |
| GP130ΔICD | Glass | 4110 | 3.90±0.46 | 11.6±1.0 |
| | MIET | 4002 | 0.84±0.05 | |
| GP130 Box1/2 | Glass | 3223 | 3.29±0.13 | 13.3±1.5 |
| | MIET | 5123 | 0.91±0.07 | |
| GP130 f-ICD + JAK1(FS) | Glass | 3382 | 3.91±0.38 | 22.1±2.0 |
| | MIET | 6956 | 1.37±0.09 | |
| GP130 f-ICD + JAK1 (+Ruxo) | Glass | 3594 | 3.85±0.09 | 22.8±2.1 |
| | MIET | 4460 | 1.48±0.12 | |
| GP130 f-ICD + JAK1 (-Ruxo) | Glass | 2925 | 3.84±0.05 | 18.4±2.2<br>22.7±2.7 |
| | MIET | 2689 | 1.25±0.13<br>1.53±0.16 | |

[1] Fluorescence lifetime and distance values are determined by Gaussian fitting of the respective histograms.

[2] Distances from silica layer.

**Supplementary Table S2: Summary of Lifetimes and Distances measured by FL-PAINT.**

| Condition | Substrate | #Localizatio | Lifetime (ns)[1] | Distance (nm)[2] |
|---|---|---|---|---|
| mEGFP-ALFAnb (in vitro) | Glass | 4760 | 3.94±0.24 | 8.2±0.8 |
| | MIET | 4165 | 0.73±0.05 | |
| GP130ΔICD | Glass | 5369 | 3.99±0.42 | 24.4±3.2 |
| | MIET | 1854 | 1.62±0.18 | |
| GP130 Box1/2 | Glass | 6627 | 3.81±0.46 | 23.0±3.7 |
| | MIET | 3509 | 1.53±0.19 | |
| GP130 f-ICD + JAK1(FS) | Glass | 4593 | 3.96±0.30 | 31.2±2.7 |
| | MIET | 5205 | 1.99±0.15 | |

[1] Fluorescence lifetime and distance values are determined by Gaussian fitting of the respective histograms.

[2] Distances from silica layer.

**Supplementary Table S3: Description of plasmids.**

| Plasmid name | Denomination | Source |
|---|---|---|
| pSems leader ALFAnb-mEGFP-TMD-HaloTag[1, 2] | mEGFP-TMD-HaloTag | This manuscript |
| pSems leader ALFAnb-gp130(K612-D645)-mEGFP[1] | GP130ΔICD | This manuscript |
| pSems leader ALFAnb-gp130(K612-D700)-mEGFP[1] | GP130 Box1/2 | This manuscript |
| pSems leader ALFAnb-gp130(K612-Q918)-mEGFP[1] | GP130 f-ICD | This manuscript |
| pSems JAK1-ΔPK/TK-mScarlet | JAK1(FS) | This manuscript |
| pSems JAK1-HaloTag | JAK1-HT | This manuscript |

[1] Leader sequence of the murine Igκ chain taken from the pDisplay vector (Invitrogen).

[2] GP130 TMD sequence: KFAQGEIEAIVVPVCLAFLLTTLLGVLFCFNKRD

**Supplementary Table S4: List of antibodies, proteins, and organic dye conjugates.**

| Antibody; protein; organic dye conjugates | Source | Catalogue number |
|---|---|---|
| Phospho-Tyrosine Mouse Monoclonal Antibody | Cell Signaling Technology | 9411 |
| Anti-pY705 STAT3 antibody (AF647 conjugate) | Cell Signaling Technology | 4324 |
| MASSIVE-TAG-X2-FAST anti-GFP | Massive Photonics | FAST-DNA-PAINT Kit |
| FAST-imager-F3-Cy3b | Massive Photonics | FAST-DNA-PAINT Kit |
| FAST-imager-F3-ATTO643 | Massive Photonics | Massive custom dyes |
| mEGFP-ALFAnb | Felker *et al.*, 2026(*1*) | N.A. |
| Anti-GFP nanobody enhancer (EN) | Wilmes *et al.*, 2020(*2*) | N.A. |
| HTL-JFX549 | Janelia Research Campus | N.A. |

**Supplementary Table S5: List of key reagents, materials, software, and suppliers.**

| Reagent | Source | Catalogue number |
|---|---|---|
| Glass coverslips | Marienfeld Laboratory glassware | 0117640 |
| Vinyltrimethoxysilane (VTMS) | Sigma Aldrich | 235768 |
| Toluene | Fisher Scientific | 14214914 |
| Poly-L-lysine-hydrochloride (PLL) | Sigma Aldrich | 2658 |
| ALFAtag | Romer Labs | Custom synthesis |
| N-Succinimidyl 13-Maleimido-11-oxo-4,7-dioxa-10-azatridecanoate (Mal-PEG2-NHS) | Tokyo Chemical Industry | M3079 |
| N-2-Hydroxyethylpiperazine-N'-2-ethane sulfonic acid (HEPES) | Carl Roth | 6763.3 |
| Dulbecco's phosphate buffered saline (PBS) | PanBiotech | P04-36500 |
| DNA-PAINT imaging buffer | Massive Photonics | FAST-DNA-PAINT Kit |
| Dulbecco's Modified Eagle's Medium (DMEM) | PanBiotech | P04-09500 |
| Fetal bovine serum (FBS) | PanBiotech | P30-3031 |
| Trypsin | Capricorn Scientific | TRY-1B |
| Linear polyethylenimine hydrochloride (PEI) | Polysciences | 24765-1 |
| NaCl | Carl Roth | 3957.1 |
| Bovine serum albumin (BSA) | Carl Roth | 8076.1 |
| Paraformaldehyde (PFA) | Sigma Aldrich | P6148 |
| Ruxolitinib (Ruxo) | Cell Signaling Technology | 83405 |
| Protease inhibitor | Serva | 39106 |
| DNase | Sigma Aldrich | DN25 |
| Lysozyme | Sigma Aldrich | L6876 |
| Isopropyl-β-D-thiogalactopyranoside (IPTG) | Thermo Fisher Scientific | R0392 |
| Imidazole | Carl Roth | 3899.4 |
| Urea | Carl Roth | 7638.1 |
| Ethylenediaminetetraaceticacid (EDTA) | Carl Roth | 8040.215 |
| OriginPro 9.0 | OriginLab | 9.0 |
| ImageJ-Fiji | Schindelin et al., 2012(*3*) | 1.54p (64 bit) |
| MATLAB R2022b | MathWorks | 2022 |
| UCSF ChimeraX | Pettersen et al., 2021(*4*) | 1.8 |
| PyMOL | Schrödinger | 3.1 |
